# Rag GTPases Suppress Renal Cystic Disease by Inhibiting TFEB Independently of mTORC1

**DOI:** 10.1101/2025.07.24.664930

**Authors:** Flaviane de Fatima Silva, Alexander R. Boucher, Huawei Li, Qingbo Chen, Maria Gaughan, Ekaterina D. Korobkina, Marie Sophie Isidor, Abigail O. Smith, Kuang Shen, Derek B. Allison, Pamela V. Tran, Gregory J. Pazour, David A. Guertin

**Author notes:** Corresponding author: David A. Guertin, PhD, Professor, Program in Molecular Medicine, UMass Chan Medical School, 366 Plantation St, N4-1099, Worcester, MA 01605. Novo Nordisk Foundation Center for Basic Metabolic Research, University of Copenhagen, Copenhagen, Denmark (Marie Sophie Isidor). Boston Combined Residency Program in Pediatrics, Boston Children’s Hospital, Boston Medical Center, Boston, MA, 02115. These authors contributed equally (Huawei Li, Qingbo Chen).

## Abstract

Aberrant mTORC1 activation in renal tubular epithelial cells (rTECs) is implicated as a critical driver of renal cystic diseases (RCDs), including autosomal dominant polycystic kidney disease (ADPKD) and tuberous sclerosis (TSC), yet its precise role remains unclear. Rag GTPases recruit mTORC1 to lysosomes, its intracellular activation site. Unexpectedly, we found that deleting *RagA/B* in rTECs, despite inhibiting mTORC1, triggers renal cystogenesis and kidney failure. We identify TFEB as the key driver of cystogenesis downstream of *RagA/B* loss and show that Rag GTPases, rather than mTORC1, are the primary suppressors of TFEB *in vivo*. We further highlight increased nuclear TFEB as a shared feature of several RCD models, whereas differences in mTORC1 activity may explain the variable efficacy of mTORC1 inhibitors. Finally, we provide evidence that nuclear TFEB, rather than mTORC1 activation, is a more consistent biomarker of cyst-lining epithelial cells in ADPKD. Overall, these findings challenge the prevailing view that mTORC1 hyperactivation is required for renal cystogenesis, which has important translational implications.

**Teaser:** A serendipitous finding uncovers the Rag GTPases as strong suppressors of renal cystogenesis with important disease implications.

## Introduction

Aberrant activation of mechanistic target of rapamycin complex 1 (mTORC1) has long been considered the key driver of renal cystic diseases (RCD), including autosomal dominant polycystic kidney disease (ADPKD) and renal cysts associated with tuberous sclerosis complex (TSC) and Birt-Hog-Dubé (BHD) syndrome (*1–5*). However, the precise mechanisms by which mTORC1 signaling drives cystogenesis across different RCDs remain poorly defined. Nevertheless, the mTORC1 inhibitor rapamycin and its analogs are under investigation as potential therapies for RCDs, including ADPKD. Yet, despite promising results in preclinical ADPKD models, human trials have shown minimal benefit for reasons that remain unclear (*6–10*), highlighting critical gaps in the understanding of how mTORC1 drives renal cystogenesis.

mTORC1 is a master regulator of cell growth and metabolism that becomes dysregulated in many human conditions, including cancer, diabetes, and aging-related pathologies. mTORC1 promotes cell growth by stimulating anabolic pathways, such as protein, lipid, and nucleic acid biosynthesis through its canonical substrates, S6K and 4E-BP1 (*11*). mTORC1 also facilitates growth by suppressing catabolic pathways like autophagy by directly phosphorylating and inhibiting the ULK1 complex (*12, 13*), and by directly phosphorylating and inhibiting Transcription Factor EB (TFEB), a transcriptional activator of autophagy, lysosomal biogenesis, and metabolic genes (*14–18*). The activation of mTORC1 occurs at lysosomes in response to both nutrients and growth factors. Mechanistically, nutrients such as amino acids stimulate mTORC1 recruitment to the lysosome via the Rag GTPase, a heterodimer composed of RagA or RagB and RagC or RagD, that is anchored to the lysosomal surface by Ragulator (*11, 19*). At the lysosome, mTORC1 is catalytically activated by Rheb, another lysosomal GTPase that is inhibited by the TSC complex and stimulated by growth factors (*20, 21*). Thus, the Rag-Regulator complex is considered the central organizer of mTORC1 activation, ensuring that mTORC1 is only activated when both sufficient nutrients and growth factors are present (*22*).

In this study, we describe an unexpected mouse model of renal cystic disease caused by the deletion of *RagA* and *RagB* in renal tubular epithelial cells (rTECs). This finding was surprising because *RagA/B* loss inhibits Rag GTPase-dependent mTORC1 activation, which is expected to suppress, rather than promote, cystogenesis. We provide evidence that TFEB activation downstream of *RagA/B* loss is sufficient to drive cystogenesis despite simultaneously inhibiting mTORC1. Our findings further reveal that Rag GTPases, rather than mTORC1 itself, serve as the primary negative regulators of TFEB in rTECs. We also find that aberrant TFEB nuclear localization—rather than aberrant mTORC1 activity—is a more consistent biomarker of the cyst-lining epithelial cells in human ADPKD samples. These results advance a new mechanistic understanding of the Rag-GTPase-mTORC1-TFEB signaling axis *in vivo* and highlight TFEB and its upstream regulatory pathways as potential therapeutic targets for renal cystic diseases, including ADPKD (*23*).

## Results

### *RagA/B* deletion in rTECs causes polycystic kidney disease

While investigating Rag GTPase function in UCP1-positive brown adipocytes, we unexpectedly discovered that *Ucp1-*Cre;*RagA/B* male mice develop kidney cysts by 4 weeks of age, which progresses to severe polycystic kidney disease by 12 weeks [Fig. S1A-C], indicated by H&E staining [Fig. S1B] and increased kidney to body weight ratio [Fig. S1C]. Cystogenesis occurred in the kidney cortex and medulla [Fig. S1D] and was associated with fibrosis [Fig. S1E], inflammation [Fig. S1F], and elevated blood urea nitrogen (BUN) levels [Fig. S1G]. A comparative analysis of 8-week-old female *Ucp1-*Cre;*RagA/B* mice also showed cyst formation with slightly less severity [Fig. S1H-K]. This was unexpected: First, because UCP1 expression is widely considered specific to brown/beige adipocytes; second, because mTORC1 activation is considered necessary for renal cystogenesis, and deleting *RagA/B* inhibits mTORC1.

To explain why *Ucp1-*Cre*;RagA/B* mice develop polycystic kidneys, we did PCR genotyping, which showed Cre-mediated recombination of the *RagA* and *RagB* floxed alleles in whole kidney samples [Fig. S2A]. Cell labeling with *Ucp1-*Cre*;mTmG* reporter mice, in which Cre induces indelible mGFP expression, pinpointed Ucp1-Cre recombination to rTECs in the medulla and some in the cortex [Fig. S2B]. Labeling with *Ucp1-*Cre*;mTmG*;*RagA/B* mice additionally localized Cre-mediated mGFP expression to epithelial cells lining the cysts [Fig. S2C]. Whole tissue Western blots from wild type and *Ucp1^-/-^*mice confirmed UCP1 protein expression in the kidney of 1-week-old mice [Fig. S2D], and UCP1 co-immunofluorescence with aquaporin 2 (AQP2) localized its expression to the collecting duct in 1-week-old mice [Fig. S2E]. These data suggested that *Ucp1-*Cre*-*mediated deletion of *RagA/B* in kidney TECs was the cause of polycystic kidney formation.

To confirm that Rag GTPase inhibition specifically in rTECs drives cystogenesis, we generated *Cadherin 16/Ksp-*Cre*;RagA/B* mice (hereafter *Ksp-*Cre*;RagA/B*), which conditionally delete *RagA/B* in distal tubular epithelial cells and collecting duct cells (*24*). Strikingly, all *Ksp-*Cre*;RagA/B* male and female mice developed renal cysts with similar trajectories [Fig. 1A-C, Fig. S3A-C]. Kidneys were visibly larger at 6 weeks in the *Ksp-Cre;RagA/B* mutants [Fig. 1A, Fig. S3A] and cyst formation was detectable by 4 weeks [Fig. 1B, Fig. S3B]. Kidney to body weight ratio substantially increased from 4 through 12 weeks in the *Ksp-*Cre*;RagA/B* mice [Fig. 1C, Fig. S1C] with cyst formation equally distributed throughout the cortex and medulla [Fig. 1D, Fig. S3D]. In males, fibrosis is detectable by 8 weeks [Fig. 1E], correlating with a robust immune response [Fig. 1F], and blood urea nitrogen (BUN) levels that are 2.9-fold higher than controls by 12 weeks [Fig. 1G] indicating kidney failure. All *Ksp-*Cre*;RagA/B* male mice have a shortened lifespan, succumbing to renal failure between 4-6 months of age [Fig. 1H]. No renal cysts or kidney failure were detected in *Ksp-* Cre*;RagA* or *Ksp-*Cre*;RagB* single knockout mice through 24 weeks, indicating compensation in suppressing renal cystogenesis [Fig. S4A-H]. Collectively, these data uncover RagA/B as potent suppressors of kidney cystogenesis.

**Figure 1.**
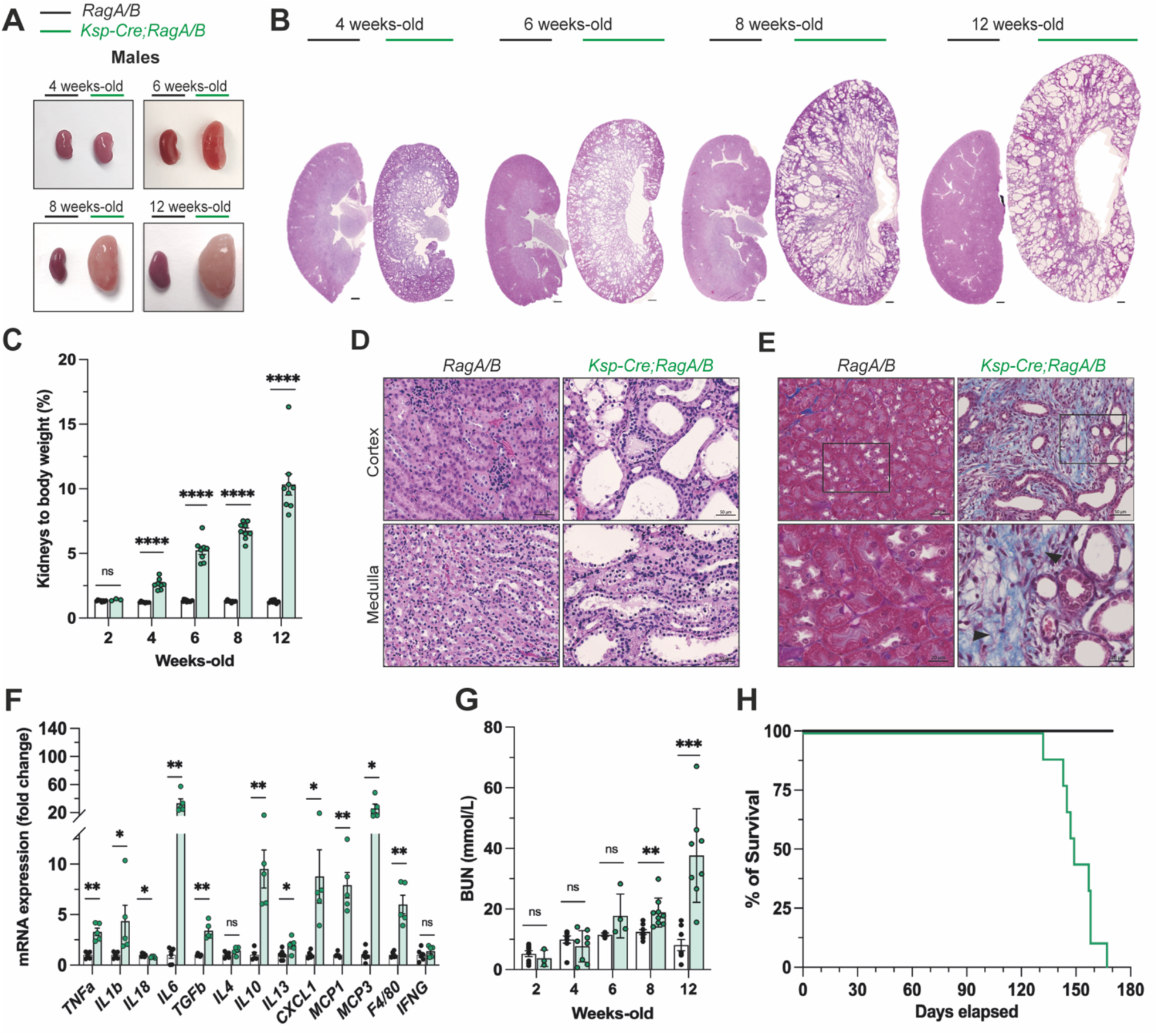
*RagA/B* deletion in kidney tubular epithelial cells promotes a striking polycystic phenotype. **(A)** Representative image from *RagA/B* floxed control and *Ksp-*Cre;*RagA/B* male mice kidney morphology and **(B)** Whole kidney H&E sections from 4 to 12 weeks of age. Scale bars = 500 µm. **(C)** Kidneys/body weight ratio from 2 to 12 weeks of age. **(D)** H&E sections showing the cyst’s location within the kidney cortex and medulla. Scale bars = 50 µm. **(E)** Low (top) and high (bottom) magnification of Mason’s Trichrome-stained kidneys showing collagen accumulation (blue) in *Ksp-* Cre;*RagA/B* at 8 weeks of age. Scale bars = 50 µm (top) and 20 µm (bottom). **(F)** Inflammation markers gene expression at 8 weeks of age. **(G)** Blood urea nitrogen (BUN) serum levels from 2 to 12 weeks of age. **(H)** Kaplan-Meier curve showing survival of *Ksp-*Cre;*RagA/B* (n = 8) relative to the floxed controls (n=8). White bars = *RagA/B* floxed control, n = 7-10; green bars = *Ksp-*Cre;*RagA/B*, n = 3-10. In **C**, **F**, and **G,** data are presented as mean ± SEM. *p<0.05, **p<0.01, and ****p<0.0001 versus the respective timepoint floxed control (unpaired Student t-test); ns = p>0.05.

### *RagA/B* loss causes cystogenesis despite inhibiting mTORC1

Aberrant mTORC1 activation is a hallmark of polycystic kidney diseases (*2–4*); thus, it was unexpected that deleting *RagA* and *RagB*—essential components of the Rag GTPase complex that activate mTORC1—induces cystogenesis. We confirmed decreased *RagA* and *RagB* mRNA and protein expression in kidneys from *Ksp-*Cre*;RagA/B* mice [Fig. 2A, 2B]. The residual expression likely reflects non-targeted cell populations present in whole kidney samples, which presents an obstacle for determining the effects of *RagA/B*-deletion on mTORC1 activity specifically in the rTECs. For example, Westerns using whole kidney tissue lysates show total protein upregulation of the mTORC1 substrates S6K and 4E-BP1, as well as AKT in the mutant kidneys [Fig. 2B], hinting at broader tissue-wide effects beyond the rTEC subpopulation.

**Figure 2.**
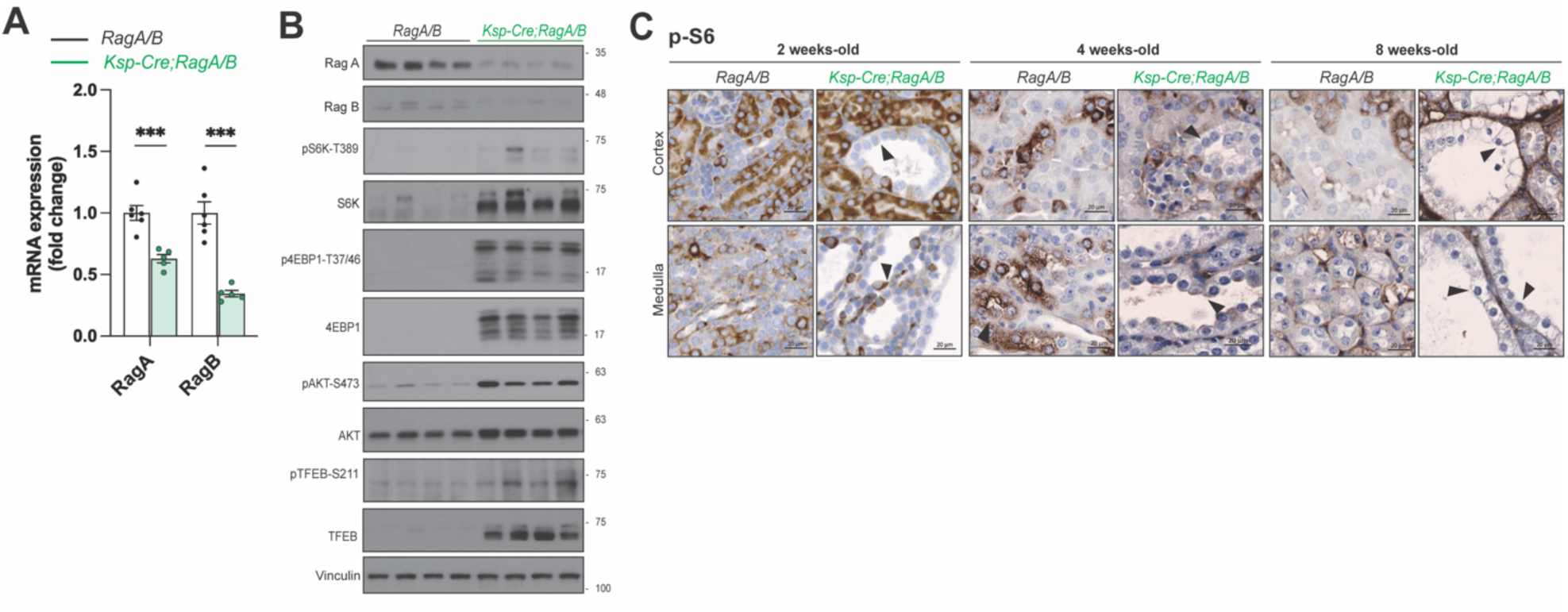
Cystogenesis is not dependent on mTORC1 activation. **(A)** *RagA* and *RagB* gene expression in kidneys of *RagA/B* floxed control and *Ksp-*Cre;*RagA/B* male mice at 6 weeks of age. **(B)** Western blotting confirmed reduced RagA and RagB protein content and mTORC1 downstream proteins. **(C)** p-S6 is low or not expressed in the cyst lining cells. IHC sections of *RagA/B* floxed control and *Ksp-*Cre;*RagA/B* male mice at 2, 4, and 8 weeks of age. Scale bars = 20 µm. White bars = *RagA/B* floxed control, n = 6; green bars = *Ksp-*Cre;*RagA/B*, n = 5. In **A,** the data are presented as mean ± SEM. ***p<0.001 versus the respective timepoint floxed control (unpaired Student *t*-test).

To circumvent this caveat and examine mTORC1 activity specifically in the rTECs that Ksp-Cre targets, we performed immunohistochemistry (IHC). Phosphorylated-S6 (p-S6), a target of S6K and a robust *in vivo* mTORC1 reporter, was absent in most of the cyst lining TECs in the medulla and cortex of *Ksp-*Cre*;RagA/B* mice at 2, 4, and 8 weeks [Fig. 2C], consistent with mTORC1 inactivation. We also detected mildly elevated p-AKT in *Ksp-* Cre*;RagA/B* rTECs at 4 and 8 weeks [Fig. S5A], which is another indicator of decreased mTORC1 signaling caused by loss of feedback inhibition of the PI3K/AKT pathway (*25–27*). Additionally, we noted a significant increase in p-S6 in the non-targeted interstitial cells, uncovering a secondary response triggered by an unknown mediator of cell crosstalk [Fig. S5B]. Nevertheless, these data indicate that renal cysts form in *Ksp-*Cre*;RagA/B* mice despite mTORC1 inhibition in rTECs.

### RagA/B suppresses TFEB activation in rTECs

Next, we investigated the effects of *RagA/B* loss on TFEB (*28–30*). Inhibition of TFEB by direct mTORC1 phosphorylation at the lysosome is considered the main mechanism of preventing TFEB nuclear localization and gene activation (*31*). It was also recently shown that TFEB is recruited to mTORC1 via a direct interaction with Rag-GTPase (*29, 32*), which is different from how canonical mTORC1 substrates like S6K and 4E-BP1, which contain a TOR-signaling (TOS) motif, are recruited to mTORC1 (*33–35*). In normal rTECs, TFEB is primarily cytosolic or distributed between cellular compartments [Fig. 3A]. Strikingly, in the *Ksp-*Cre*;RagA/B* mice, TFEB is concentrated in the nucleus of nearly all cyst-lining TECs during early cyst formation (2 weeks), and it becomes more prominent in the nucleus as the disease progresses (4 and 8 weeks) [Fig. 3A], which correlates with an increase in total TFEB protein levels in whole kidney lysates [Fig. 2B].

**Figure 3.**
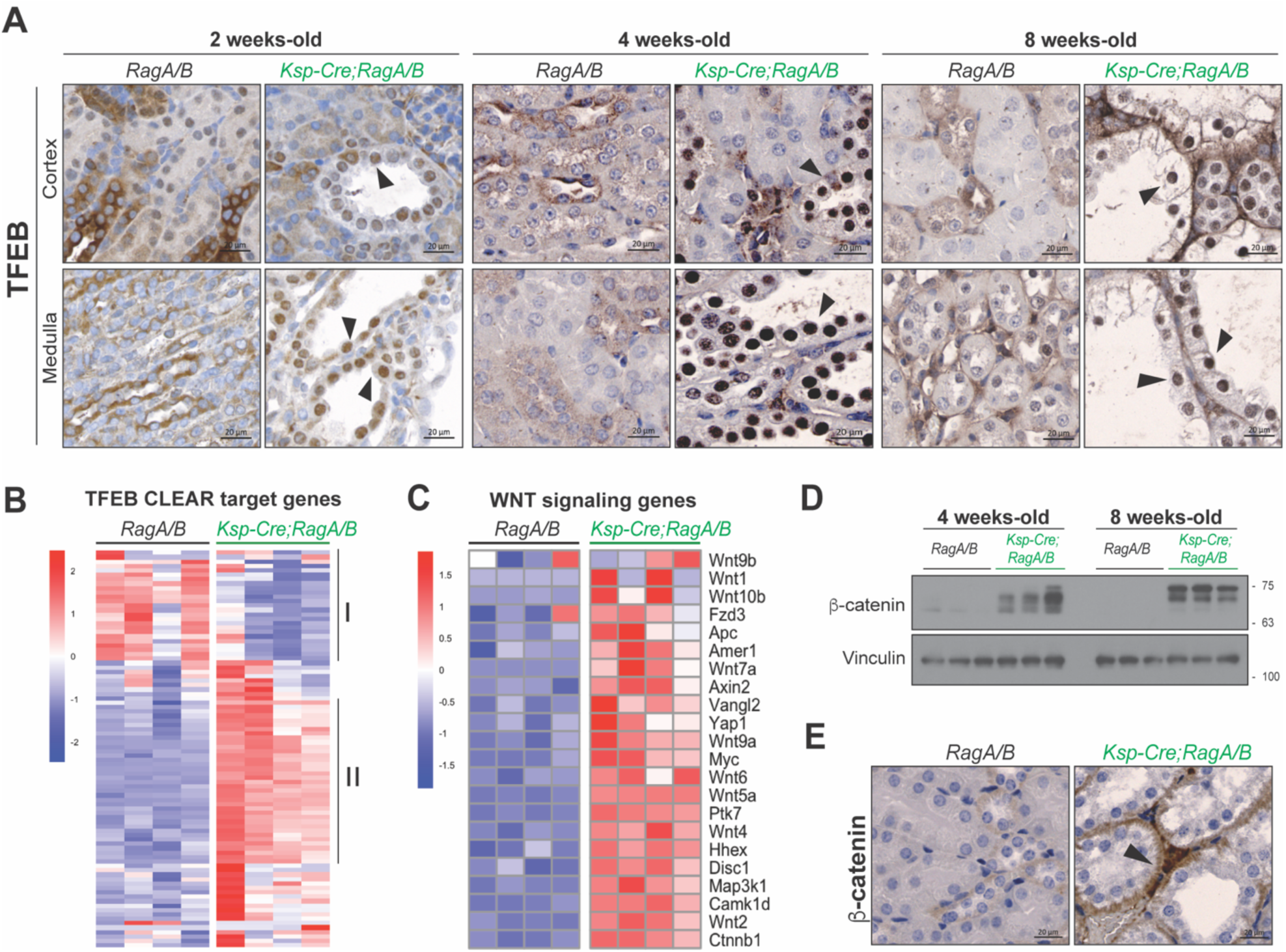
TFEB is the driver of kidney cystogenesis in the absence of *RagA/B*. **(A)** TFEB localization throughout cyst progression at 2, 4, and 8 weeks of age. **(B)** Heatmaps portraying TFEB CLEAR target gene set expression differences (Log2FoldChange) between *RagA/B* floxed controls and *Ksp-* Cre;*RagA/B* male mice at 6 weeks of age. **(C)** WNT signaling gene set expression differences (Log2FoldChange) between *RagA/B* floxed controls and *Ksp-*Cre;*RagA/B* male mice at 6 weeks of age. **(D)** Whole kidney lysate western blotting for **β**-catenin at 4 and 8 weeks of age. **(E) β**-catenin IHC sections of *RagA/B* floxed control and *Ksp-*Cre;*RagA/B* male mice at 8 weeks of age. Scale bars = 20 µm.

Consistent with TFEB activation in *Ksp-*Cre*;RagA/B* mutant kidneys, bulk RNA-sequencing shows upregulation of direct TFEB target genes, called Coordinated Lysosomal Expression and Regulation (CLEAR) genes, in the cystic kidneys (*36*) [Fig. 3B], noting that a subset of CLEAR genes also decreases [Fig. 3B], indicated by group I and II genes, respectively. Analysis of the broader TFEB target gene network (also called the CLEAR network) showed a similar pattern [Fig. S6A]. We also observe increased WNT/β-catenin signaling, which has been associated with increased TFEB activation (*37*), indicated by an increase in WNT target gene expression in cystic kidneys [Fig. 3C], increased β-catenin expression in whole tissue lysates [Fig. 3D], and β-catenin localization to adherens junctions in the mutant rTECs [Fig. 3E]. These data suggest that RagA/B is a key suppressor of TFEB and WNT activation *in vivo*.

### Reduced efficacy of rapamycin in *Ksp*-Cre;*RagA/B* mice compared to *Ksp*-Cre;*Tsc2* mice

Several preclinical studies show that rapamycin reduces cyst formation across multiple models of RCD (*3*). Rapamycin is particularly effective at reducing cystic burden in *Ksp-* Cre*;Tsc2* kidneys, a model of renal cystogenesis in tuberous sclerosis associated with aberrant activation of both mTORC1 and TFEB (*3*). In this model, rapamycin completely relieved cystic burden and corrected both aberrant mTORC1 activation and TFEB nuclear localization, leading to a model in which high mTORC1 activity promotes TFEB nuclear localization by an unknown mechanism to drive cystogenesis (*3*). We compared our whole kidney bulk RNA-seq data from the *Ksp-*Cre*;RagA/B* mice to published bulk RNA-seq data from the kidneys of age-matched *Ksp-*Cre*;Tsc2* mice (*3*), which showed remarkable similarity [Fig. 4A]. The similarity in gene signatures was surprising given that mTORC1 is inhibited by RagA/B loss while mTORC1 is activated by TSC2 loss; nevertheless, the similarity suggests at least partial overlapping molecular pathology.

**Figure 4.**
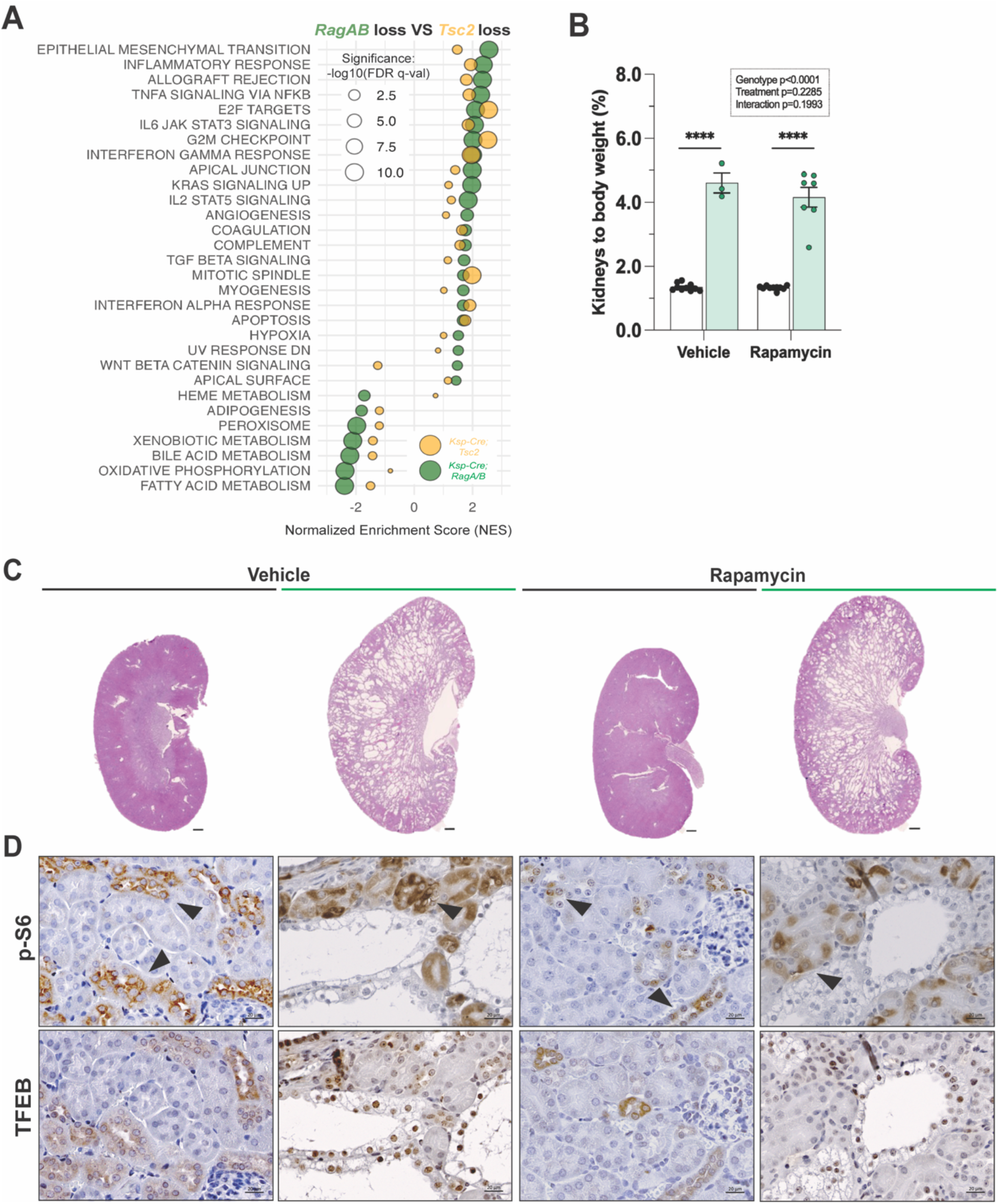
Reduced efficacy of rapamycin in *Ksp*-Cre;*RagA/B* mice compared to *Ksp*-Cre;*Tsc2* mice. **(A)** GSEA comparison of molecular hallmarks from whole kidney bulk-RNAseq of *Ksp-*Cre;*RagA/B* and *Ksp-*Cre;*Tsc2* kidneys (only includes hallmarks with FDR q-val </= 0.05 in the *Ksp-*Cre;*RagA/B* dataset). **(B)** Kidneys/body weight ratio at P50 (floxed control + vehicle, n = 10; *Ksp-Cre;RagA/B* + vehicle, n = 3; floxed control + rapamycin, n = 10; *Ksp-*Cre;*RagA/B* + vehicle, n = 7). **(C)** Representative whole kidney sections from *RagA/B* floxed control and *Ksp-*Cre;*RagA/B* treated with vehicle or rapamycin 1.0 mg/Kg at P50. Scale bars = 500 µm. **(D)** p-S6 and TFEB IHC kidney sections from *RagA/B* floxed control and *Ksp-*Cre;*RagA/B* male mice treated with vehicle or Rapamycin 1.0 mg/kg. Scale bars = 20 µm. White bars = *RagA/B* floxed control; green bars = *Ksp-* Cre;*RagA/B*. In **B,** data are presented as mean ± SEM. ****p<0.0001 versus the respective treated floxed control–genotype effect (Two-Way ANOVA).

This raised the question of whether rapamycin might be effective at reducing cystic burden in the *Ksp-*Cre*;RagA/B* mice. We hypothesized that rapamycin would be less effective in the *Ksp-*Cre*;RagA/B* model because mTORC1 is already inhibited by RagA/B loss. To test this, we treated *Ksp-*Cre*;RagA/B* and littermates floxed controls with Rapamycin (1.0 mg/Kg) from P25 to P50 [Fig. 4B] following the exact protocol previously shown to be effective at reverting kidney cysts in *Ksp-*Cre*;Tsc2* mice (*3*). Consistent with our hypothesis, rapamycin offered no benefit in reducing kidney size or cystogenesis in the *Ksp-*Cre*;RagA/B* mice when delivered using the same dose and treatment regimen that was highly effective at reducing cystic burden in *Ksp-*Cre*;Tsc2* mice [Fig. 4C, 4D]. By IHC, we observed that this rapamycin treatment regimen reduced the pS6 signal detected in the interstitial cells of *Ksp-*Cre*;Tsc2* mice [Fig. 4D], but did not affect TFEB nuclear localization, which remained nuclear [Fig. 4D]. This is consistent with cystogenesis in both *Ksp*-Cre;*RagA/B* and *Ksp*-Cre;*Tsc2* mice being driven by distinct mechanisms that converge on TFEB activation, and it reveals that not all preclinical models of RCD are necessarily driven by aberrant mTORC1 activation and thus may not respond equally to rapamycin treatment.

### *Tfeb* is required for cyst formation downstream of *RagA/B* loss

In other models of renal cystogenesis, aberrant TFEB/TFE3 is thought to promote cystogenesis by activating mTORC1 in a feedback activation loop (*3, 5, 38*). Therefore, we next asked whether TFEB was required for cystogenesis downstream of *RagA/B* loss, in which mTORC1 is inhibited. For this, we generated *Ksp-*Cre*;RagA/B;Tfeb* triple knockout (TKO) mice. Strikingly, at 6 weeks of age, there was no difference in kidney size between control and TKO mice [Fig. 5A, 5B], no cysts were detected throughout the TKO kidneys [Fig. 5C], and there were no signs of kidney dysfunction based on BUN levels, which were similar to *wild-type* control [Fig. 5D]. IHC confirmed TFEB loss in rTECs, especially in the medulla, where collecting ducts and loop of Henle are located [Fig. 5E]. Using the TFEB target genes *Atg9b* and *Ctsd* as TFEB activity markers, we confirmed that *Tfeb* deletion normalizes TFEB target gene expression [Fig. 5F]. *Tfeb* deletion also normalized total S6K, 4E-BP1, and AKT protein levels in whole kidney lysates [Fig. 5G]. We noted that pAKT remained modestly elevated in mutant whole kidney lysates, which is consistent with mTORC1 inhibition [Fig. 5G], which we confirmed by IHC for pS6 and pAKT, respectively [Fig. 5H, 5I]. Thus, TFEB is required for cyst formation caused by *RagA/*B loss.

**Figure 5.**
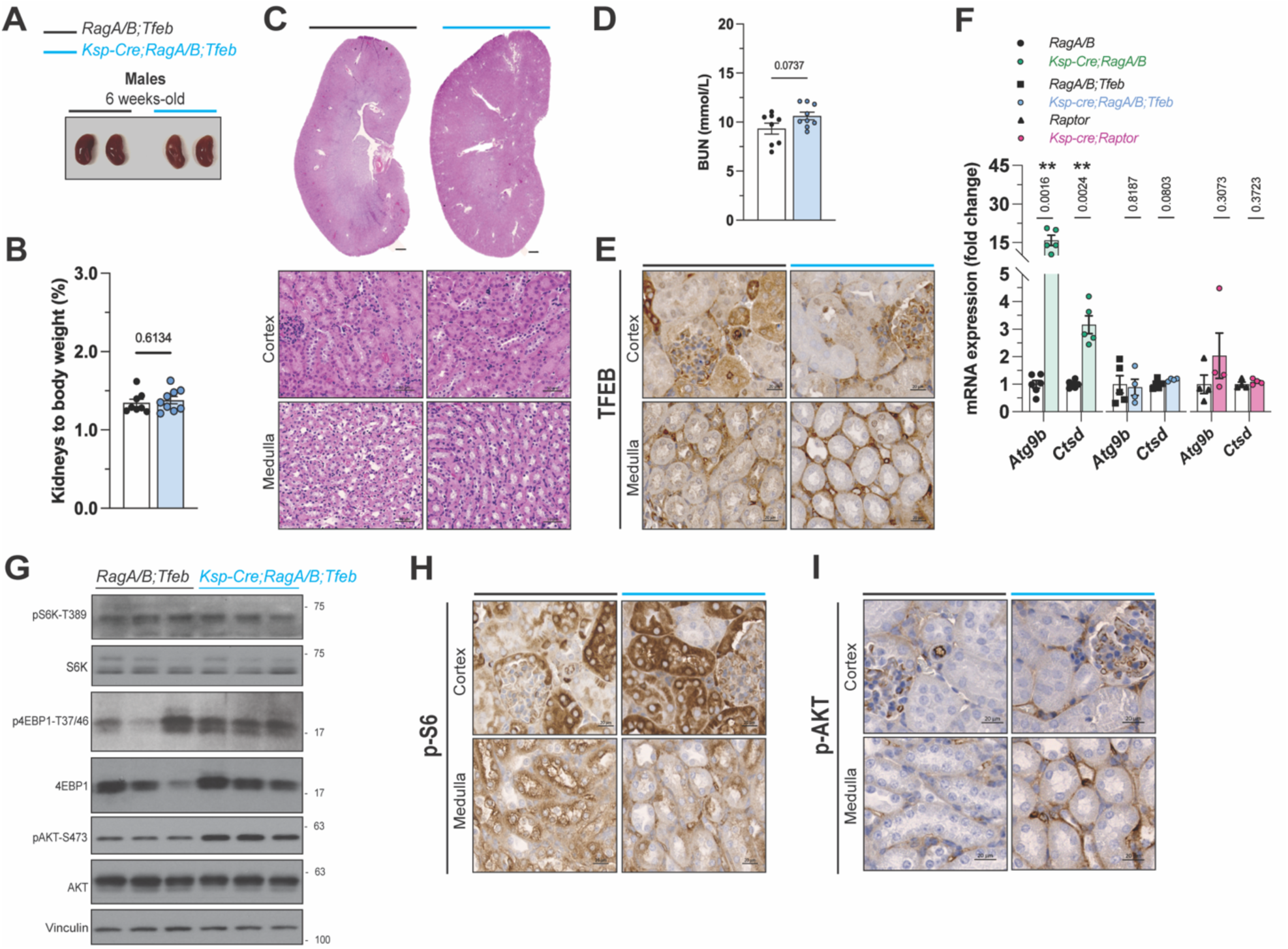
*Tfeb* is required for cyst formation downstream of *RagA/B* loss. **(A)** Representative image from *RagA/B;Tfeb* floxed control and *Ksp-*Cre;*RagA/B;Tfeb* male mice kidney morphology at 6 weeks of age. **(B)** Kidneys/body weight ratio (*RagA/B;Tfeb* floxed control, n = 8; *Ksp-*Cre*;RagA/B;Tfeb*, n = 9). **(C)** Kidney H&E sections showing absence of cysts. Scale bars = 500 µm (whole kidney), 50 µm (cortex and medulla sections). **(D)** Blood urea nitrogen (BUN) serum from *RagA/B;Tfe*b floxed control and *Ksp-*Cre;*RagA/B;Tfeb* levels at 6 weeks of age. **(E)** TFEB IHC sections of *RagA/B;Tfeb* floxed control and *Ksp-*Cre;*RagA/B;Tfeb* male mice at 6 weeks of age. Scale bars = 20 µm. **(F)** Expression of TFEB target genes *Atg9b* and *Ctsd* (*RagA/B* floxed control, n = 6; *Ksp*-Cre;*RagA/B* n = 5; *RagA/B;Tfeb* floxed control, n = 5; *Ksp*-Cre;*RagA/B;Tfeb*, n = 4; *Raptor* floxed control, n = 5; *Ksp-*Cre;*Raptor*, n = 4). **(G)** Whole kidney lysate western blotting for mTORC1 downstream proteins. **(H) p**-S6 IHC sections of *RagA/B;Tfeb* floxed control and *Ksp-*Cre;*RagA/B;Tfeb* male mice at 6 weeks of age. Scale bars = 20 µm. **(I) p**-AKT IHC sections of *RagA/B;Tfeb* floxed control and *Ksp-*Cre;*RagA/B;Tfeb* male mice at 6 weeks of age. Scale bars = 20 µm. White bars = *RagA/B, RagA/B;Tfeb* or *Raptor* floxed controls; green bars = *Ksp-*Cre;*RagA/B*; blue bars = *Ksp-*Cre;*RagA/B;Tfeb*; pink bars = *Ksp-*Cre;*Raptor*. In **B**, **D**, and **F,** data are presented as mean ± SEM. p values versus the respective floxed control (unpaired Student’s *t*-test).

### Rag GTPase, not mTORC1, is the main TFEB suppressor *in vivo*

mTORC1 is widely considered the major negative regulator of TFEB activity (*15, 31, 39*). However, *Ksp-*Cre*;Raptor* mice, which are predicted to both inhibit mTORC1 and trigger nuclear TFEB, just like the *Ksp-*Cre*;RagA/*B mice, have visibly smaller kidneys with no cysts and normal BUN levels [Fig. 6A-D] (*40*). As expected, deleting *Raptor* both decreased pS6 and increased pAKT in rTECs, respectively [Fig. 6E, 6F]. However, TFEB did not accumulate in the nucleus of *Raptor* mutant rTECs [Fig. 6G], and the mRNA *Atg9b* and *Ctsd* were not elevated by *Raptor* loss [Fig. 5F]. Notably, the total protein levels of S6K and 4E-BP1 were elevated in *Ksp-*Cre*;Raptor* mutant whole kidneys [Fig. 6H], like in the *Ksp-*Cre*;RagA/*B mice [Fig. 2B], suggesting this phenomenon is more linked to impaired Rag-GTPase and/or mTORC1 activity but not TFEB activation. Overall, the observation that both *Ksp-* Cre*;RagA/B* and *Ksp-*Cre*;Raptor* models reduce mTORC1 signaling, but only *RagA/B* deletion activates TFEB, reinforcing TFEB activation as the critical driver of cystogenesis and reveals that, *in vivo*, Rag GTPases—not mTORC1—serve as the primary negative regulators of TFEB.

**Figure 6.**
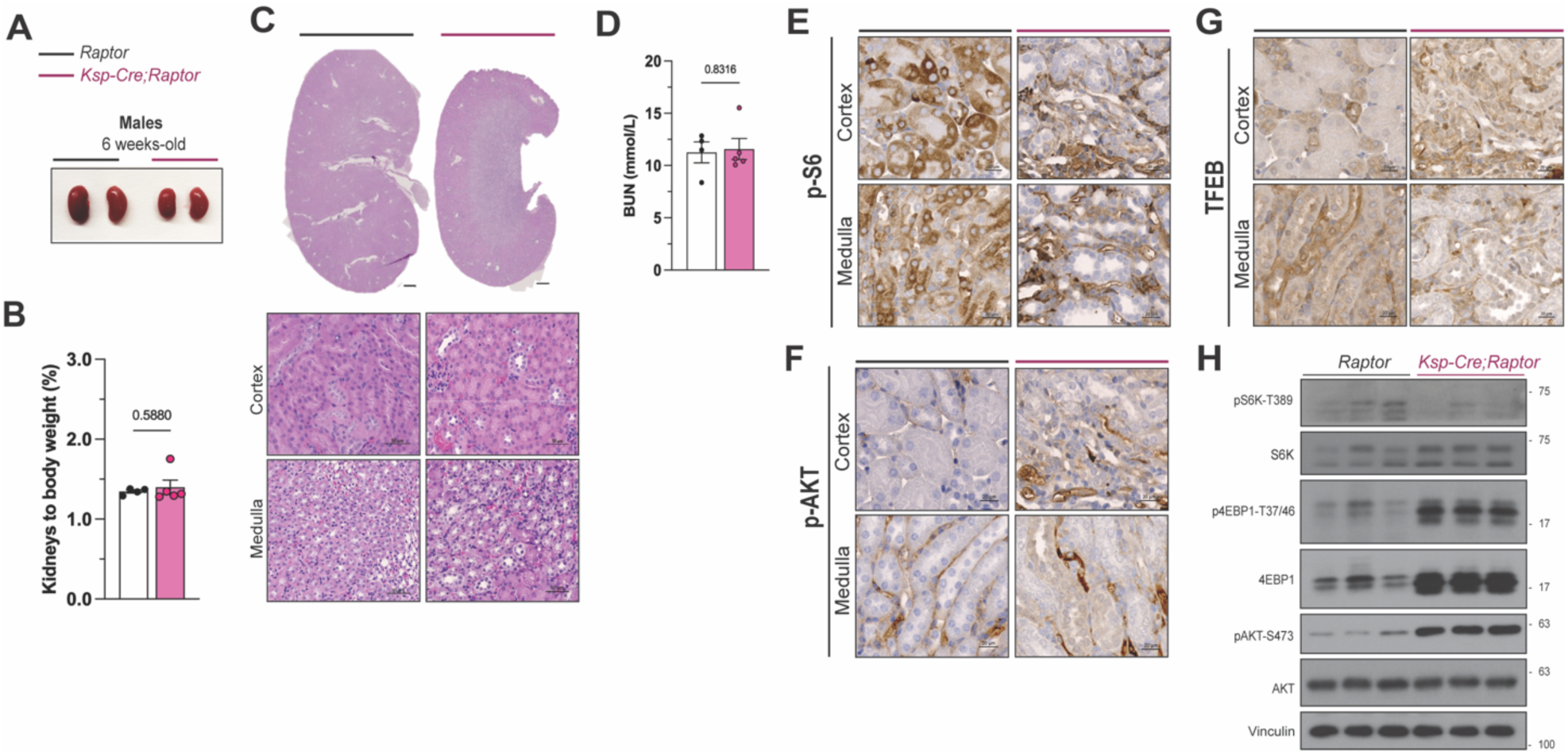
Rag GTPase, not mTORC1, is the main TFEB suppressor *in vivo.* **(A)** Representative image from *Raptor* floxed control and *Ksp-*Cre;*Raptor* male mice kidney morphology at 6 weeks of age. **(B)** Kidneys/body weight ratio and **(C)** H&E sections showing absence of cysts. Scale bars = 500 µm (whole kidney), 50 µm (cortex and medulla sections). **(D)** Blood urea nitrogen (BUN) serum from *Raptor* floxed control and *Ksp-*Cre;*Raptor* levels at 6 weeks of age. **(E)** p-S6 IHC, **(F)** p-AKT, and **(G)** TFEB IHC sections of *Raptor* floxed control and *Ksp-*Cre;*Raptor* male mice at 6 weeks of age. Scale bars = 20 µm. **(H)** Whole kidney lysate western blotting for mTORC1 downstream proteins. White bars = *Raptor* floxed control, n = 4; pink bars = *Ksp-*Cre;*Raptor,* n = 5. In **B** and **D,** data are presented as mean ± SEM. p values versus the floxed control (unpaired Student’s *t*-test).

### Nuclear TFEB is more consistent than pS6 as a biomarker in ADPKD

We also compared bulk RNA-seq data from *Ksp*-Cre;*RagA/B* cystic kidneys to previously published RNA-seq data from kidneys of postnatal day 7 (*P7*) *HoxB7*-Cre;*Pkd1* and *HoxB7*-Cre;*Pkd2* mice (*41*), established ADPKD models with reasonably well-matched bulk RNA-seq data. This analysis also revealed remarkable similarity in gene signatures between the *RagA/B* mutant kidneys and both the *Pkd1* and *Pkd2* mutant gene signatures [Fig. 7A]. Epithelial-mesenchymal transition (EMT), E2F targets, IL6/JACK-STAT signaling, TGFβ signaling, and WNT/β-catenin signaling were among the commonly upregulated pathways with established links to ADPKD (*42*). We also generated a PKD-hallmark gene set based on published genes associated with ADPKD models [Fig. 7B]. Most of the genes we identified showed directional changes in the *RagA/B* mutant kidneys that are consistent with ADPKD, including increases in *Tacstd2* and *Gprc5a* expression (*43*). This overlap between *Ksp-*Cre*;RagA/B* kidneys and models of ADPKD, in addition to TSC, suggests that deregulation of TFEB is a shared feature of different renal cystic diseases, as recently proposed (*2*). A recent single-cell RNA-sequencing study using human ADPKD samples also identified TFEB target genes as being significantly upregulated (*44*).

**Figure 7.**
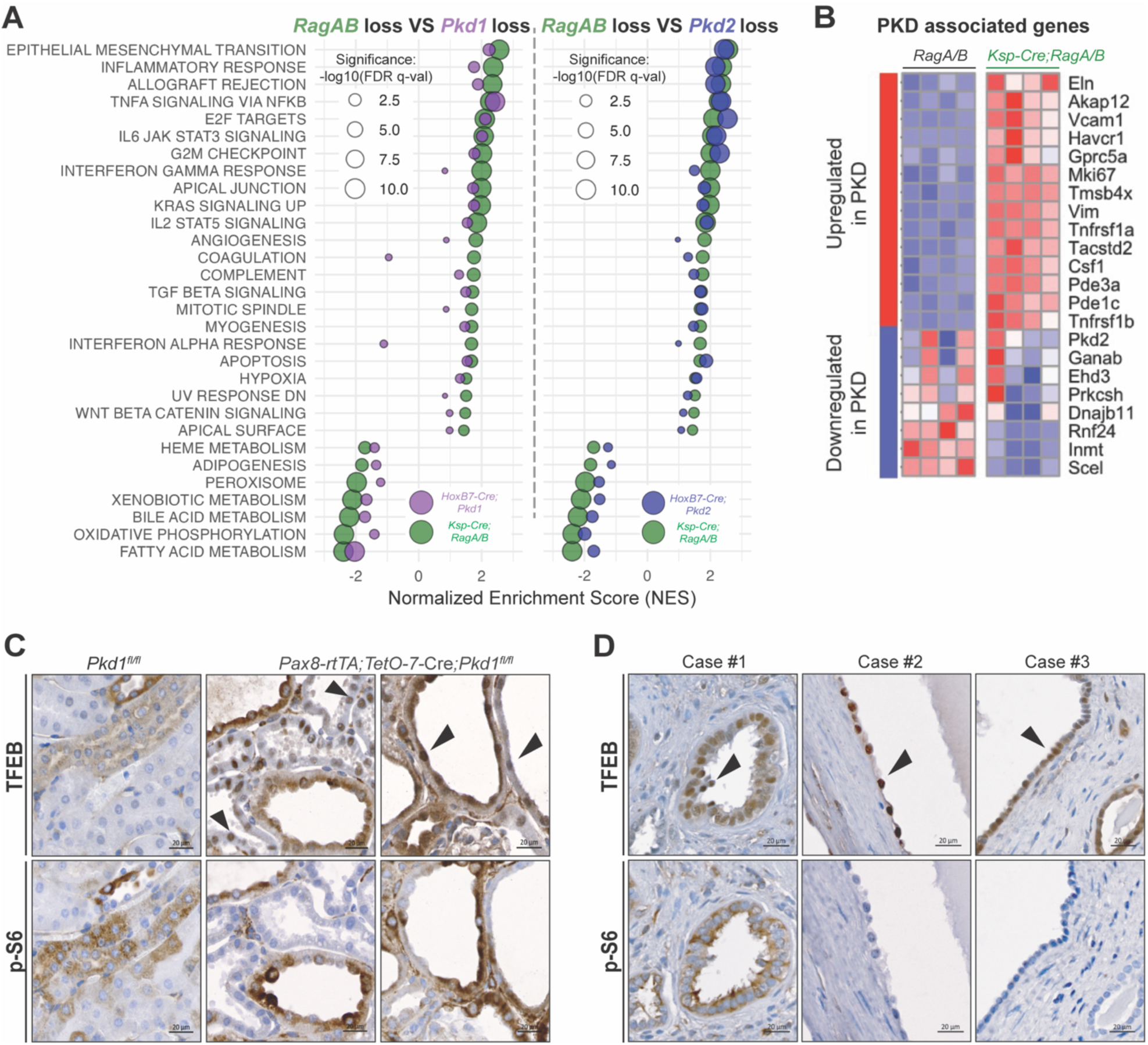
RagA/B deletion overlaps with ADPKD models, with TFEB activation as a consistent biomarker. **(A)** GSEA comparison of molecular hallmarks from whole kidney bulk-RNAseq of *Ksp-* Cre;*RagA/B* and *HoxB7-Cre;Pkd1* (left) or *HoxB7-*Cre;*Pkd2* (right) kidneys (only includes hallmarks with FDR q-val </= 0.05 in the *Ksp-Cre;RagA/B* dataset). **(B)** Heatmaps portraying PKD-associated genes expression differences (Log2FoldChange) between *RagA/B* floxed controls and *Ksp*-Cre;*RagA/B* male mice at 6 weeks of age. **(C)** Representative TFEB and p-S6 IHC from *Pkd1*^fl/fl^ and *Pax8-rTTA;Tet-O-7-*Cre;*Pkd1^fl/fl^* male mice and **(D)** kidney sections from ADPKD patients. Scale bars = 20 µm.

To investigate this further in ADPKD, we compared TFEB nuclear localization to pS6 as biomarkers of renal cystic cells in the *Pax8-rtTA;Tet-O-7-*Cre*;Pkd1* model, a widely accepted preclinical mouse model of ADPKD (*44, 45*). This revealed a heterogeneous labeling pattern; for example, some cystic cells showed strong nuclear TFEB signal but little to no pS6 [Fig. 7C]—a pattern consistent with the *Ksp-*Cre*;RagA/B* model. Other cyst-lining cells showed elevated total TFEB protein that was distributed between both the nucleus and cytosol, which also had elevated pS6 [Fig. 7C]. These data hint at temporal and/or spatial differences in how mouse rTECs may reprogram mTORC1 and TFEB activity in response to *Pkd1* loss. Nevertheless, it confirms aberrant TFEB expression as a prominent biomarker in a pre-clinical model of ADPKD.

Finally, we examined three human ADPKD kidney sections from two patients with PKD1 mutations and a third patient with a family history of *PKD1* mutations. In all three cases, aberrant TFEB nuclear localization was consistently present in the cyst-lining epithelial cells [Fig. 7D]. However, the pS6 signal was inconsistent. For example, in the patient 1 samples, we observed high nuclear TFEB that overlapped with high pS6; but in patient 2 and 3 samples, we observed nuclear TFEB but no pS6, which resembles the pattern we observed in the *Ksp-*Cre*;RagA/B* model. Thus, aberrant TFEB activation may be a more consistent biomarker of late-stage ADPKD than aberrant mTORC1 activation, warranting a deeper investigation into the function of the Rag-GTPase/TFEB pathway in ADPKD.

## Discussion

Aberrant mTORC1 activation is widely considered a hallmark driver of renal cystic diseases (RCDs), including ADPKD and TSC. However, its precise molecular role in cyst formation and whether it functions similarly across different types of RCDs has remained unclear. In this study, we discovered that RagA and RagB, which are required for RagC and RagD activity and mediate mTORC1 activation at lysosomes (*22*), function as redundant suppressors of renal cystogenesis. This demonstrates that the conventional Rag GTPase-activated mTORC1 pathway is not essential for renal cystogenesis. Instead, we show that TFEB drives cyst formation downstream of Rag GTPase loss, independent of hyperactivation of the canonical Rag GTPase-mTORC1 pathway. These findings highlight the broad relevance of TFEB—both functionally and as a biomarker—in RCDs and underscore key gaps in understanding if and how to target the mTORC1 pathway across different RCD subtypes.

Another key finding is that Rag GTPases, rather than mTORC1, serve as the primary negative regulators of TFEB in rTECs *in vivo*. This was uncovered by showing that Raptor loss, unlike RagA/B loss, results in minimal changes in TFEB localization *in vivo*. This was surprising because the prevailing model, based largely on *in vitro* studies, is that mTORC1-dependent phosphorylation of TFEB is the primary mechanism restricting its nuclear localization and activity (*31*). Recent structural data indicate that RagA and RagC directly interact with TFEB, in addition to mTORC1, to facilitate their interaction at lysosomes (*29*), which is different than how S6K and 4E-BP1 interact with mTORC1 (*46*). However, our genetic data suggest that direct interaction between Rag GTPase and TFEB may be the dominant mechanism restraining TFEB nuclear localization *in vivo*, with mTORC1-dependent phosphorylation being an auxiliary mechanism to fine-tune TFEB activity. *In vitro*, siRNA knockdown of *Raptor* in human retinal pigment epithelial cells induced TFEB nuclear localization (*30*), seemingly at odds with our *in vivo* findings. However, that study also reported that *Raptor* knockdown caused a substantial fraction of TFEB to redistribute to lysosomes. One possibility is that Raptor loss frees up TFEB binding sites at lysosomes, and *in vivo*—within a physiologically relevant context—this lysosomal sequestration may be sufficient to restrain pathological TFEB activation, even when mTORC1-dependent TFEB phosphorylation is inhibited. Another possibility is that a compensatory kinase or other mechanism may substitute for mTORC1 in phosphorylating and inhibiting TFEB upon *Raptor* loss. Future biochemical and genetic studies will be needed to test these hypotheses.

We also show that the cystic kidneys in *Ksp-*Cre*;RagA/B* mutants show considerable molecular overlap with models of renal cystogenesis in TSC (*Tsc2* loss) and ADPKD (both *Pkd1* and *Pkd2* loss), including aberrant TFEB activation (*2, 3, 41*), suggesting TFEB activation is a common denominator across several RCDs, while mTORC1’s role may be more variable. Genetic studies in mice support a critical role for TFEB in TSC (*3, 32*), and in Birt-Hogg-Dubé Syndrome, another rare genetic disorder associated with increased risk of kidney cancer (*5, 47*). A functional role for TFEB in ADPKD remains elusive. Nevertheless, these findings substantiate the potential value of targeting TFEB or its downstream effectors in RCDs. It is interesting that a rapamycin treatment regimen highly effective at reverting cysts in a *Ksp-*Cre*;Tsc2* model of renal cystogenesis, also driven by TFEB activation, was ineffective in the *Ksp-*Cre*;RagA/B* model. This aligns with our finding that TFEB, but not canonical mTORC1 signaling, is essential for cystogenesis in *Ksp-*Cre*;RagA/B* mice and may hint at different pathological mechanisms that converge upon TFEB activation.

A note on UCP1 in the kidney. The discovery that RagA/B suppresses renal cystogenesis was serendipitous, arising from an unexpected phenotype following *Ucp1*-Cre– mediated deletion of *RagA* and *RagB*. Although UCP1 is classically regarded as specific to brown and beige adipocytes, multiple studies have hinted at its expression in non-adipocyte cell types, including the kidney (*48*). However, this has been debated (*49*). Here, using multiple protein-based assays and a *Ucp1* knockout control mouse, we demonstrate UCP1 expression mainly in renal TECs of the collecting duct, explaining why *Ucp1-Cre;RagA/B* developed renal cysts, and challenging the long-standing assumption that UCP1 is restricted to adipose tissues. UCP1 expression in the kidney may have been overlooked in other studies due to dynamic regulation, restricted expression to only a small subset of cells, and/or technical limitations of RNA sequencing, particularly when its mRNA expression is low. While a potential function for UCP1 in renal epithelial cells remains unclear, it may relate to a role in mitigating mitochondrial stress (*50*). Notably, human UCP1 polymorphisms (e.g., 3826 A/G) have been linked to hypertension independent of obesity (*51, 52*), raising the possibility of a previously overlooked role in kidney physiology. Future studies are warranted to define whether UCP1 functions in this context.

One limitation of our study is that it does not rule out the possibility that unidentified mTORC1 substrates that might be regulated in a Rag GTPase and/or lysosome-independent manner may contribute to cyst formation in *RagA/B*-deleted rTECs. Mounting evidence suggests that Rag GTPase–independent mTORC1 signaling can occur, including *in vivo*; however, a consensus model and defined functional role have yet to emerge (*53–55*). Higher dosing of rapamycin and genetic ablation of Raptor in *Ksp-*Cre*;RagA/B* would help address this possibility. Overall, our findings that canonical mTORC1 signaling is not essential for renal cystogenesis underscore the critical need to further investigate the relative contributions of mTORC1, TFEB, and Rag GTPases across RCDs, including their temporal and spatial deregulation, to develop optimal treatments.

## Materials and Methods

### Animal studies

B6.Cg-Tg(Cdh16-cre)91Igr/J (strain #012237, hereafter named *Ksp-Cre*), C57BL/6J mice (stock #000664, hereafter named *wild type*), B6.129-Ucp1tm1Kz/J (Strain #:003124, hereafter named *Ucp1^-/-^*), and *R26R-mTmG* (stock #007676) were purchased from The Jackson Laboratories. *Ucp1-Cre* mice (strain #024670) were provided by E. Rosen. *Tfeb^fl/fl^* mice were provided by E. Henske. *Ksp-*Cre*;RagA, Ksp-*Cre*;RagB*, *Ksp-* Cre*;RagA/B, Ksp-* Cre*;RagA/B;Tfeb, Ksp-*Cre*;Raptor, Ucp1-*Cre*;RagA/B, Ucp1-* Cre*;Raptor, Ucp1-*Cre*;mTmG*, and *Ucp1-*Cre*;RagA;RagB;mTmG* mice were generated by crossing in our facility. Mice were housed at 22 °C, in 50% ± 20% humidity, and a 12-h light–dark cycle, with free access to water and standard chow diet (Prolab Isopro RMH 3000). All experiments were approved by the UMass Chan Medical School IACUC (Protocol #202100035) and conducted with adherence to the National Institutes of Health Guide for the Care and Use of Laboratory Animals.

Treatment with rapamycin 1.0 mg/kg was performed as previously described [28], with 11 injections from P25 to P50. Paired control groups received vehicle (0.1% ethanol in sterile 0.9% NaCl). By the timepoints described (2 to 12 weeks old, or P50), the body weight was assessed, and euthanasia was performed by exsanguination (cardiac puncture under 2.0 % Isoflurane anesthesia), followed by cervical dislocation and kidney dissection. The euthanasia for all experiments was conducted at the same time of the day to minimize possible circadian variations, and all mice were in a fed state. The weight of the two kidneys was normalized and expressed as a percentage of body weight (Kidneys to body weight (%)). Blood samples were centrifuged for 10 minutes at 4 °C and 10.000 rpm, and the serum was aliquoted and kept at −80 °C until further analysis. Tissues were snap frozen using a liquid nitrogen-cooled Wollenburg clamp and transferred into 2-ml microcentrifuge tubes with a pre-cooled 5-mm metal bead. Tissues were homogenized using CryoMill (Retsch) at 25 Hz for 2 min. Tissue powder was weighed out into pre-cooled microcentrifuge tubes to be used in the analysis described below.

### Tissue histology and Immunohistochemistry

Whole kidneys from all genotypes described were sectioned in the coronal plane and fixed in 10% formalin in PBS. Embedding, sectioning, and staining (H&E, Mason’s Trichrome, and immunohistochemistry - IHC) were performed by the UMCMS Morphology Core facility. Paraffin-embedded kidney blocks from *Pkd1^fl/fl^* and *Pax8-rtTA;Tet-O-7*-Cre;*Pkd1^fl/fl^* mice were provided by the PKD Rodent Model Core of the Kansas PKD Research and Translational Core Center at KUMC (U54 DK126126) and the PKD Research Resource Consortium (PKD RRC). Kidney sections from ADPKD patients were provided by Derek B. Allison (University of Kentucky College of Medicine IRB # 79922; Table S1). For IHC, the sections were deparaffinized, rehydrated, submitted to antigen retrieval (10 mM Sodium Citrate buffer, pH 6.0), blocking of endogenous peroxidase with 3% hydrogen peroxide solution, and then blocked with 5% goat serum. The slides were incubated overnight (4 °C) with the primary antibodies p-S6-S240/244 (1:1000, 5364S, Cell Signaling), TFEB (1:100, 13372-1-AP, Proteintech), or p-Akt-S473 (1:200, 4058L, Cell Signaling). The sections were then washed and incubated with HRP-conjugated Goat anti-Rabbit IgG secondary antibody (1:200, AS014, Abclonal), developed with SignalStain® DAB Substrate (8059P, Cell Signaling), and mounted. Full slide scans were taken at 20X using a Zeiss Axio-Scan.Z1. Zeiss Zen Blue 3.7 was used for image acquisition and processing.

### Immunofluorescence

Kidneys and BAT from *Ucp1-*Cre*;mTmG* and *Ucp1-*Cre*;RagA/B;mTmG* mice were processed as previously described (*56*). The slides containing frozen sections were washed in PBS and mounted with Fluoromount-G with DAPI (00-4959-52, Invitrogen). For UCP1 and Aquaporin 2 (AQP2) co-staining, the kidney sections were deparaffinized, rehydrated, submitted to antigen retrieval with 10 mM Sodium Citrate buffer pH 6.0 at 120°C for 30 min, and blocked with 4% goat serum in 0.05% TBST with 0.1 % fish skin gelatin and 0.1% Triton-X100. Primary antibodies UCP1 (1:500, ab10983, Abcam) and aquaporin 2 – AQP2 (1:100, SAB5200110, Sigma), diluted in 0.05% TBST with 0.1% fish skin gelatin were added overnight at 4 °C. The slides were incubated with the secondary antibodies for 1 h at room temperature (Alexa Fluor 488 - A11008, 1:1000, and Alexa Fluor 555-A-21428, 1:1000, Invitrogen). The slides were mounted with Fluoromount-G with DAPI. All immunofluorescence images were taken at 10X or 20X magnification using a Zeiss Confocal LSM 900+ Airyscan Microscope, and Zeiss Zen Blue 3.7 was used for image processing.

### Blood Urea Nitrogen assessment

The serum was diluted 1:1 with sterile 0.9% NaCl, and the urea was measured following the urease method by a commercial kit (E-BC-K183-S, Elabscience). The assay sensitivity is 0.114 mmol/L, inter-assay CV: 4.7%, and intra-assay CV: 4.6%. The Blood urea nitrogen (BUN) final concentration was expressed as mmol/L.

### Gene expression analysis

For qPCR, RNA was isolated from tissues using Qiazol (#79306, QIAGEN)-chloroform and processed through RNeasy kit (#74106, QIAGEN). RNA (1.0 μg) was reverse-transcribed to complementary DNA using a High-Capacity cDNA Reverse Transcription kit (#4368813, Applied Biosystems). Quantitative PCR with reverse transcription (qRT–PCR) was performed in 10-μl reactions using SYBR Green PCR master mix (CW0955, CWBio).

The reaction was performed using StepOnePlus (Applied Biosystems), and melting curves were run on every plate for all genes to ensure efficiency and specificity. TATA-box binding protein (*Tbp*), acidic ribosomal phosphoprotein P0 (*36b4*), and beta (β)-actin (*Actb*) gene expression geometric mean was used for normalization of each gene. The primer sequences used are listed in Table S2.

### RNAseq

RNA-Seq analyses were performed on the ViaFoundry platform66 by Novogene. Briefly, sequence reads were processed with cutadapt (v4.4) to remove adapters and trim low-quality bases. Gene expression was quantified using salmon (v1.9.0) against the mouse genome assembly mm10 Gencode m25 basic gene annotation. Differential gene expression analyses were performed using DESeq2 (v1.40.2). Genes failing a minimum read count threshold (10 reads for each replicate in at least one condition) were removed before differential expression analysis. Gene set enrichment analysis was performed on preranked DESeq2-normalized RNA-seq data using GSEA Desktop v4.3.3 (build 16; Broad Institute, 4 Feb 2024) against the MSigDB Hallmark collection with 1,000 weighted gene-set permutations. All plots were generated using R packages ggplot2 (v.3.4.2), ggrepel (v0.9.3), and pheatmap (v1.0.12) or GraphPadPrism10.

### Immunoblot analysis

Cryomilled samples from kidneys were lysed in RIPA buffer (150 mM NaCl, 50 mM HEPES at pH 7.4, 0.1% SDS, 1% Triton X-100, 1% glycerol, 2 mM EDTA, and 0.5% deoxycholate) containing protease and phosphatase inhibitor cocktail. Protein lysates were mixed with 5× SDS sample buffer, boiled at 95 °C, separated by SDS–PAGE, transferred to a polyvinylidene difluoride (PVDF) membrane, and subjected to immunoblot analysis. The primary and secondary antibodies and the respective dilutions used are listed in Table S3.

### Statistical Analysis

Data were tested for distribution (Shapiro–Wilk test) and homogeneity of variances (F test). For comparisons between each genotype described and its respective floxed controls, unpaired Student t-test or Two-Way ANOVA, as indicated in the figure captions. In the t-test, when the F test was significant, the Welch correction was applied. The data are presented as mean ± standard error of mean (SEM), and the level of significance assumed was 5% (p < 0.05). All analyses were performed using GraphPad Prism 10.2.

## Acknowledgments

We thank David M. Sabatini and Alejo Efeyan for providing the *RagA/B* mice strain, Andrea Ballabio and Elizabeth P. Henske for the *Tfeb* mice strain, Kansas PKD Research and Translational Core (U54DK126126), and the Polycystic Kidney Disease Research Resource Consortium (PKD RRC) for the *Pkd1* mouse model FFPE kidney sections.

## Author contributions

Conceptualization: FFS, ARB, GJP, DAG

Methodology: FFS, ARB, AS, DBA, PT, GJP, DAG

Investigation: FFS, ARB, HL, QC, MG, EK, MSI, AS, DBA, PT.

Data Analysis: FFS, ARB

Supervision: GJP, DAG

Writing—original draft: FFS, AB, GJP, DAG

Writing—review & editing: FFS, ARB, HL, QC, MG, EK, MSI, AS, KS, DBA, PT, GJP, DAG.

## Data and materials availability

All data needed to evaluate the conclusions in the paper are present in the paper and/or the Supplementary Materials.

