## Supplemental Data for "Rag GTPases Suppress Renal Cystic Disease by Inhibiting TFEB Independently of mTORC1"

**This PDF file includes:**

Figs. S1 to S6
Tables S1 to S3

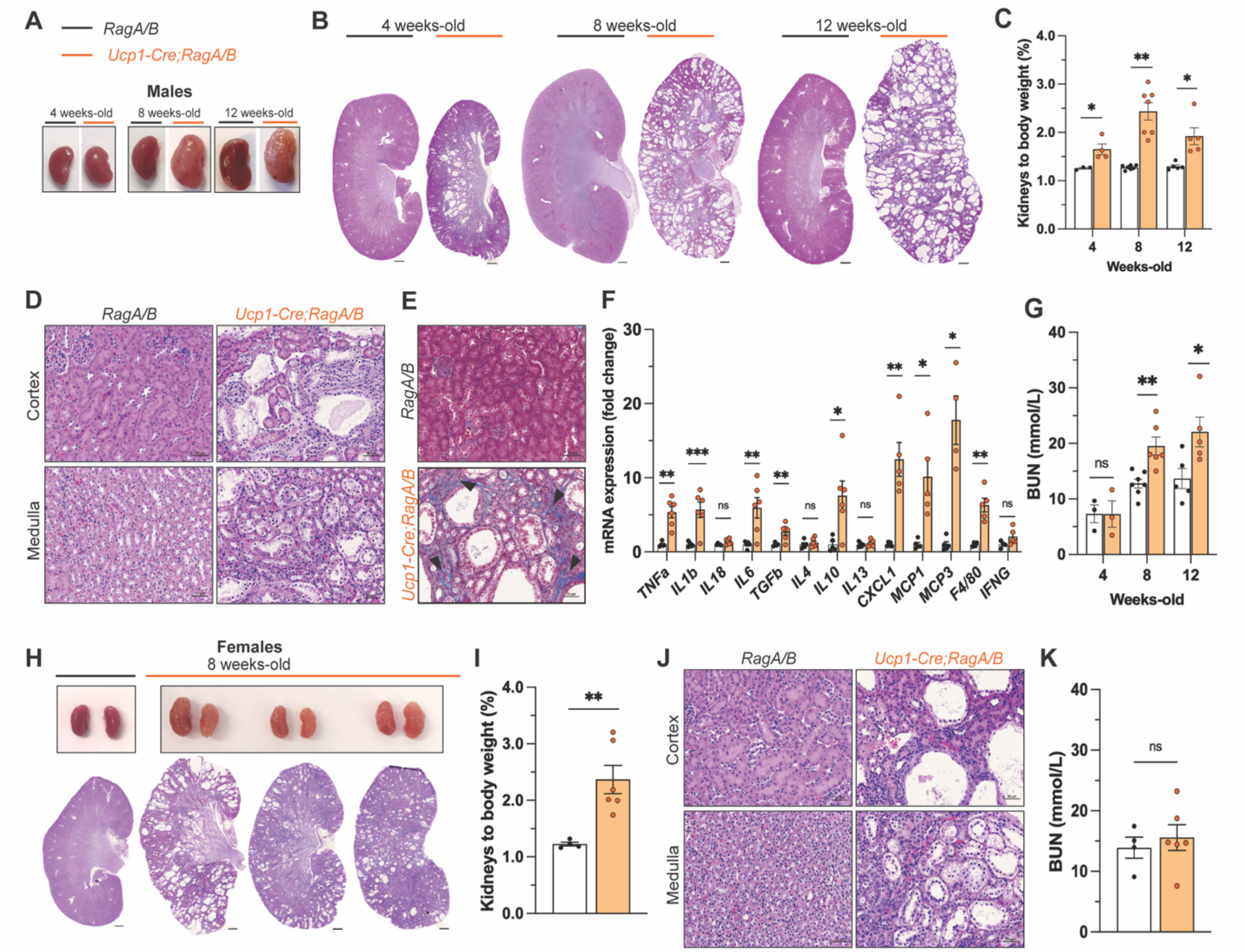


**Fig. S1. Unexpected polycystic kidney phenotype in *Ucp1*-Cre*;RagA/B* mice.**
**(A)** Representative image from *RagA/B* floxed control and *Ucp1*-Cre*;RagA/B* male mice kidney morphology from 4 to 12 weeks of age. **(B)** Whole kidney H&E sections from 4 to 12 weeks of age. Scale bars = 500 µm. **(C)** Kidneys/body weight ratio from 4 to 12 weeks of age (floxed control, n = 3-8; *Ucp1*-Cre*;RagA/B*, n = 4-7). **(D)** H&E kidney sections showing cyst's location within cortex and medulla in male mice. Scale bars = 50 µm. **(E)** Mason's Trichrome stained kidneys showing collagen accumulation (blue) in *Ucp1-Cre;RagA/B* male mice at 8 weeks of age. Scale bars = 50 µm. **(F)** Inflammation markers in male mice kidneys at 8 weeks of age (floxed control, n = 6; *Ucp1*-Cre*;RagA/B*, n = 4-6). **(G)** Blood urea nitrogen (BUN) serum levels of male mice from 4 to 12 weeks of age (floxed control, n = 3-7; *Ucp1*-Cre*;RagA/B*, n = 3-6). **(H)** Representative image from RagA/B floxed control and *Ucp1*-Cre*;RagA/B* female mice kidney morphology and whole kidney H&E sections at 8 weeks of age. **(I)** Female mice kidneys/body weight ratio at 8 weeks of age (floxed control, n = 4; *Ucp1*-Cre*;RagA/B*, n = 6). **(J)** H&E kidney sections showing cyst's location within cortex and medulla in female mice. Scale bars = 50 µm. **(K)** Blood urea nitrogen (BUN) serum levels of female mice at 8 weeks of age (floxed control, n = 4; *Ucp1*-Cre*;RagA/B*, n = 6). White bars = *RagA/B* floxed control; orange bars = *Ucp1*-Cre*;RagA/B*. In **C**, **F**, **G**, **I**, and **K** data is presented as mean ± SEM. *p<0.05, **p<0.01, and ***p<0.001 versus the respective timepoint floxed control (unpaired Student *t*-test); ns = p>0.05.

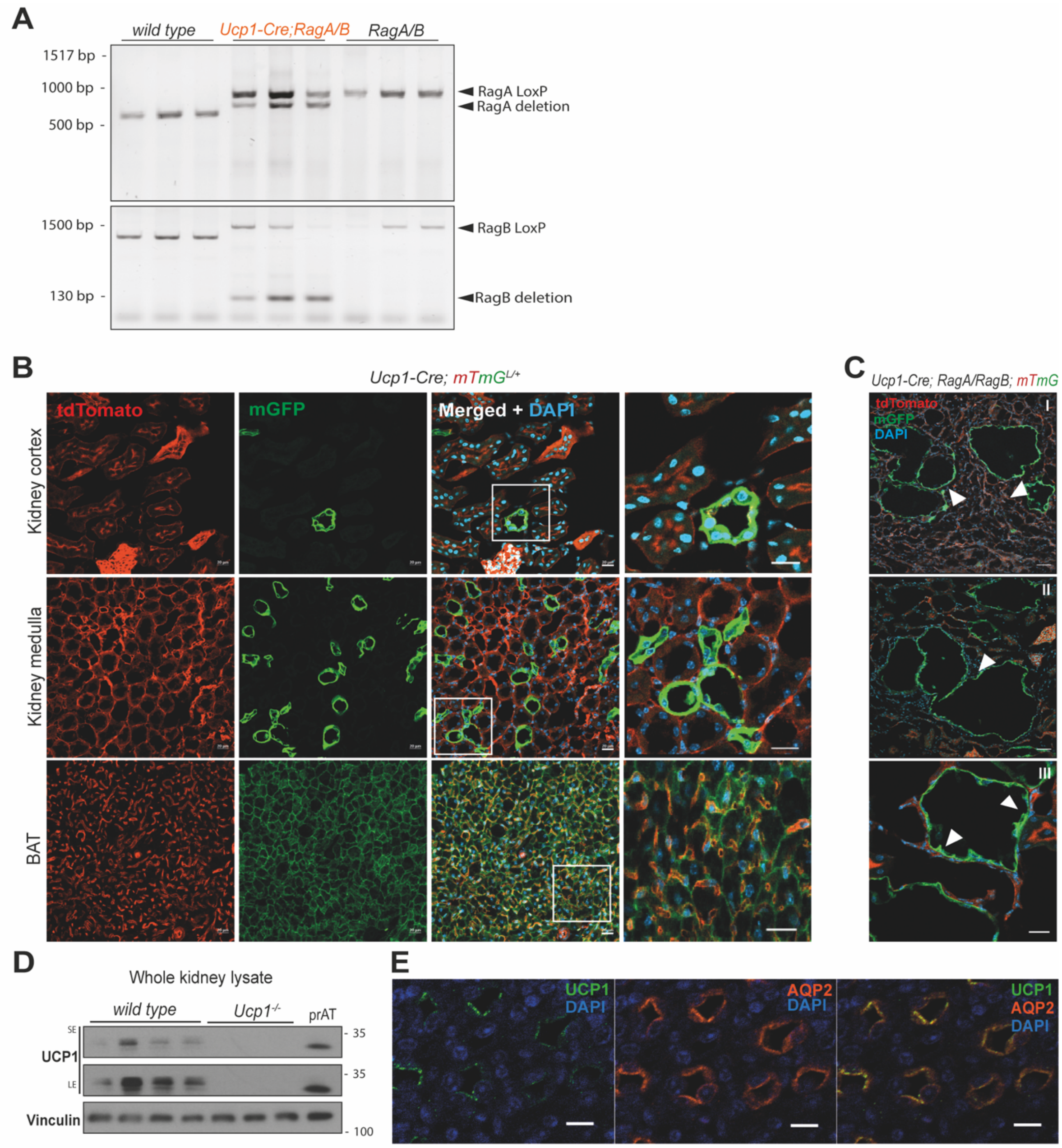


**Fig. S2. UCP1 is expressed in kidney tubular epithelial cells. (A)** DNA gel confirming kidney *RagA* and *RagB* allele deletion. **(B)** Representative kidney sections from *Ucp1*-Cre*;mTmG* male mice showing Cre-positive (mGFP) cells in cortex and medulla. Brown adipose tissue (BAT) was used as a positive control for Ucp1-Cre labeling. Scale bars = 20 µm. **(C)** Kidney sections from *Ucp1*-Cre*;RagA/B;mTmG* male mice showing cyst-lining epithelial cells as Cre-positive (mGFP). White arrows highlighting the cyst epithelial cells. Scale bars = 50 µm (I, II), and 20 µm (III). **(D)** Western blot of whole kidney lysates from wild type and *Ucp1*$^{-/-}$ male mice at 1 week of age. Perirenal adipose tissue (prAT) was used as positive control for UCP1 expression. SE = short exposure; LE = long exposure time. **(E)** Immunofluorescence showing co-localization of UCP1 (green) and Aquaporin 2 (AQP2; orange) in collecting duct epithelial cells. DAPI was used as a nuclear marker. Scale bars = 10 µm.

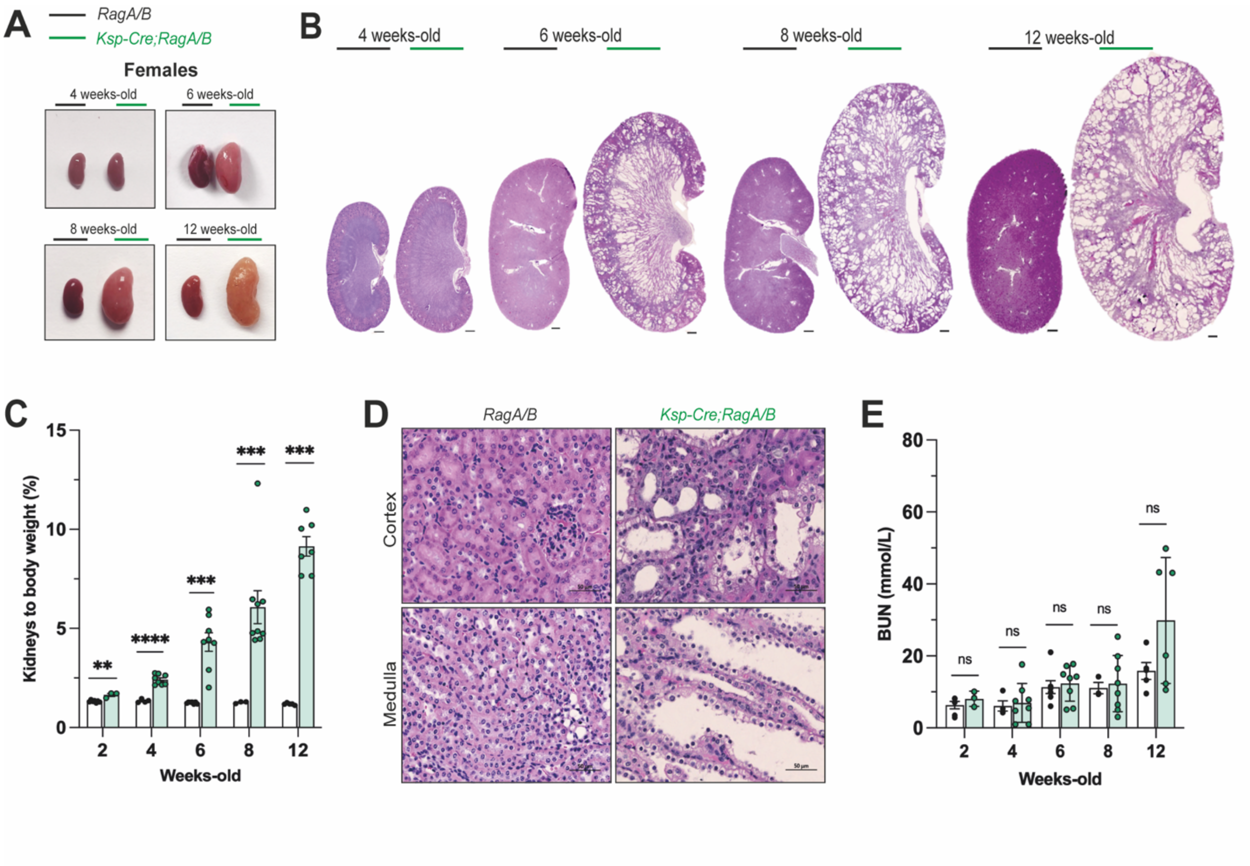


**Fig. S3. *Ksp*-Cre;*RagA/B* female mice share the same cystic phenotype to males.**
**(A)** Representative image from *RagA/B* floxed control and *Ksp*-Cre;*RagA/B* female mice showing kidney morphology from 4 to 12 weeks of age. **(B)** Whole kidney H&E sections from 4 to 12 weeks of age. Scale bars = 500 µm. **(C)** Kidneys/body weight ratio from 2 to 12 weeks of age. **(D)** H&E sections showing cyst location within kidney cortex and medulla of female mice. Scale bars = 50 µm. **(E)** Blood urea nitrogen (BUN) serum levels from 4 to 12 weeks of age. White bars = *RagA/B* floxed control, n = 4-6; green bars = *Ksp*-Cre;*RagA/B*, n = 6-8). In **C** and **E** data is presented as mean ± SEM. ***p<0.001, and ****p<0.0001 versus the respective timepoint floxed control (unpaired Student t-test); ns = p>0.05.

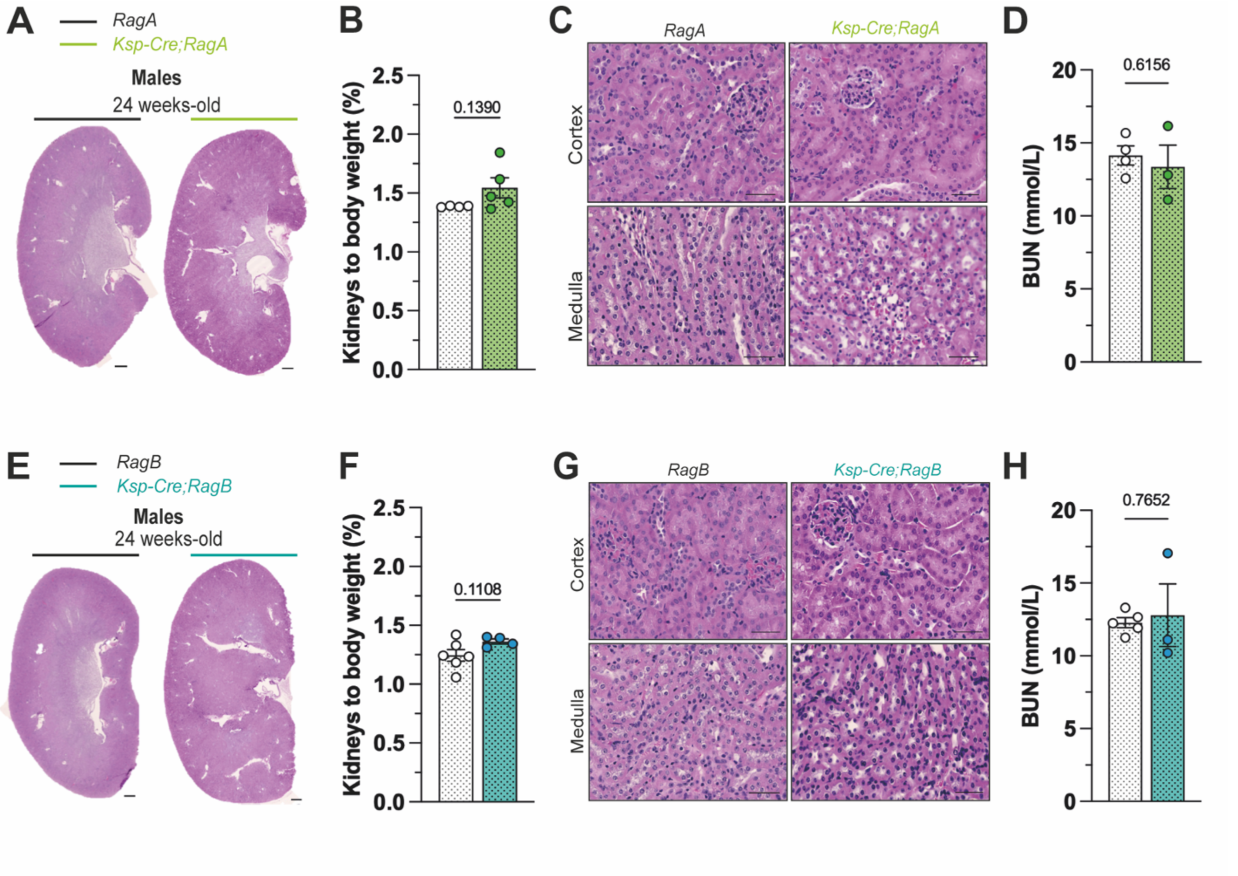


**Fig. S4. Single *RagA* or *RagB* deletion does not promote cyst development. (A)** Whole kidney H&E sections from *RagA* floxed control and *Ksp*-Cre;*RagA* male mice at 24 weeks of age. Scale bars = 500 µm. **(B)** Kidneys/body weight ratio at 24 weeks of age. **(C)** Representative *RagA* floxed control and *Ksp*-Cre;*RagA* H&E sections of kidney cortex and medulla. Scale bars = 50 µm. **(D)** Blood urea nitrogen (BUN) serum levels at 24 weeks of age. **(E)** Whole kidney H&E sections from *RagB* floxed control and *Ksp*-Cre;*RagB* male mice at 24 weeks of age. Scale bars = 500 µm. **(F)** Kidneys/body weight ratio at 24 weeks of age. **(G)** Representative *RagB* floxed control and *Ksp*-Cre;*RagB* H&E sections of kidney cortex and medulla. Scale bars = 50 µm. **(H)** Blood urea nitrogen (BUN) serum levels at 24 weeks of age.White dotted bars = *RagA* floxed control (n = 4) or *RagB* floxed control (n = 6); green dotted bars = *Ksp*-Cre;*RagA* (n = 3-5); turquoise dotted bars = *Ksp*-Cre;*RagB* (n = 3-4). In **B**, **D**, **F**, and **H** data is presented as mean ± SEM. p values versus the respective floxed control (unpaired Student *t*-test).

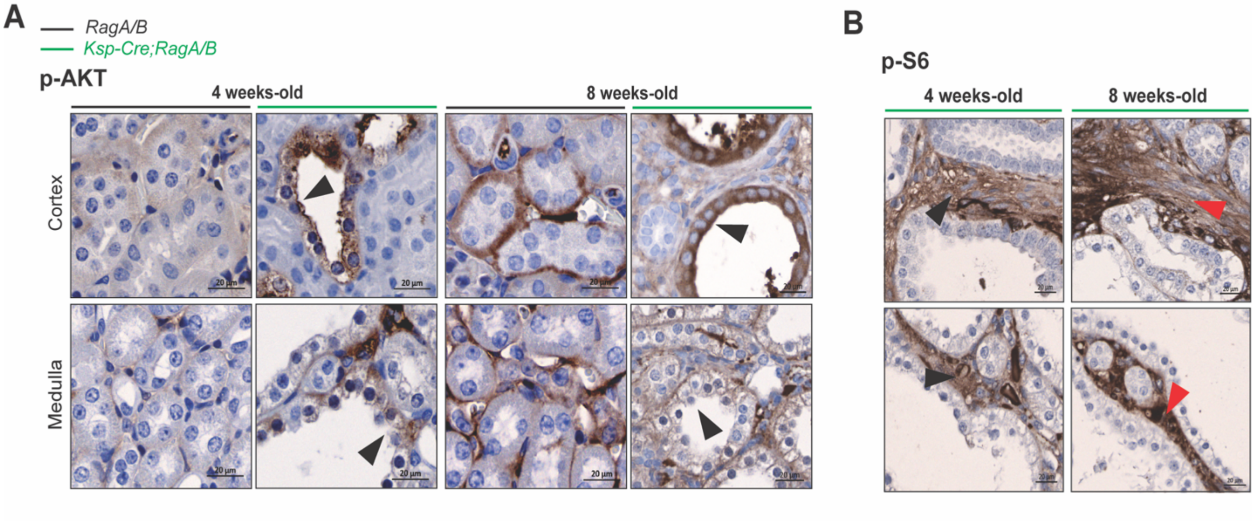


**Fig. S5. p-AKT labeling confirms mTORC1 inhibition in cyst-lining TECs, while interstitial cells show mTORC1 hyperactivation. (A)** p-AKT IHC sections of *RagA/B* floxed control and *Ksp*-Cre;*RagA/B* male mice at 4 and 8 weeks of age (black arrows). **(B)** Representative IHC sections of *Ksp*-Cre;*RagA/B* male mice at 4 and 8 weeks of age evidencing high p-S6 (red arrows) in the interstitial cells. Scale bars = 20 µm.

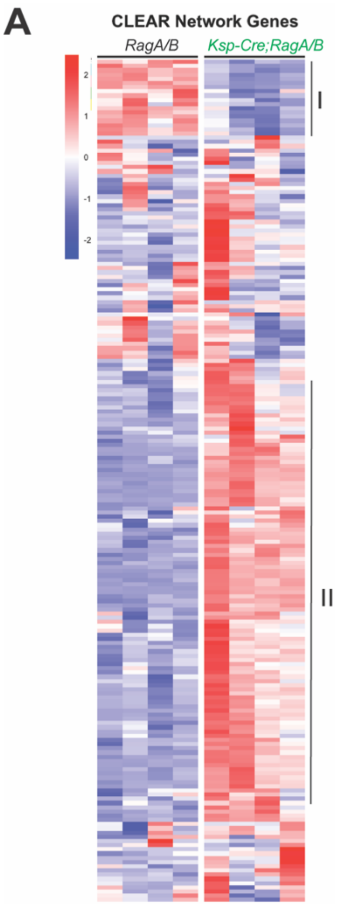


**Fig. S6. (A)** Heatmaps portraying TFEB CLEAR Network gene set expression differences (Log2FoldChange) between *RagA/B* floxed controls and *Ksp*-Cre;*RagA/B* male mice at 6 weeks of age.

| Case # | Clinical ADPKD/Mutation | Age at Surgery | Sex | BMI | Associated Comorbidities |
|---|---|---|---|---|---|
| 1 | Yes*, *Pkd1* | 44 | F | 30 | Hyperlipidemia, hypertension |
| 2 | Yes*, unknown | 48 | M | 35 | T2DM, hyperlipidemia, hypertension, secondary polycythemia |
| 3 | Yes*, *Pkd1* c.4551C>A | 50 | F | 29 | T2DM, hypertension |

* Family history consistent.

**Table S1. Details of the human ADPKD samples used in this study.**

| Gene | | Sequence |
|---|---|---|
| *36b4 (acidic ribosomal phosphoprotein P0)* | **F** | GCGACCTGGAAGTCCAACTAC |
| | **R** | ACGTTGTCTGCTCCCACAAT |
| *Actb (β-actin)* | **F** | ATGGAATCCTGTGGCATCCA |
| | **R** | CGCTCAGGAGGAGCAATGAT |
| *Atg9b (Autophagy-related 9B)* | **F** | AGCTATCATCAGCGGAATGG |
| | **R** | GCGAAGGAGGAAGGTTGTAA |
| *Ctsd (Cathepsin D)* | **F** | GCTTCCGGTCTTTGACAACCT |
| | **R** | CACCAAGCATTAGTTCTCCTCC |
| *Cxcl1 (C-X-C Motif Chemokine Ligand 1)* | **F** | CTGGGATTCACCTCAAGAACATC |
| | **R** | CAGGGTCAAGGCAAGCCTC |
| *F4/80 (Fibroblast growth factor 4 receptor)* | **F** | CTTTGGCTATGGGCTTCCAGTC |
| | **R** | GCAAGGAGGACAGAGTTTATCGTG |
| *IFNg (Interferon gamma)* | **F** | ACAGCAAGGCGAAAAAGGATG |
| | **R** | TGGTGGACCACTCGGATGA |
| *IL-10 (Interleukin 10)* | **F** | CTTACTGACTGGCATGAGGATCA |
| | **R** | GCAGCTCTAGGAGCATGTGG |
| *IL-13 (Interleukin 13)* | **F** | CCTGGCTCTTGCTTGCCTT |
| | **R** | GGTCTTGTGTGATGTTGCTCA |
| *IL-18 (Interleukin 18)* | **F** | ACAGGCCTGACATCTTCTGC |
| | **R** | TCTGACATGGCAGCCATTGT |
| *IL-1b (Interleukin 1 beta)* | **F** | GGTGTGTGACGTTCCCATTA |
| | **R** | TCCTGACCACTGTTGTTTCC |
| *IL-4 (Interleukin 4)* | **F** | GGTCTCAACCCCCAGCTAGT |
| | **R** | GCCGATGATCTCTCTCAAGTGAT |
| *IL-6 (Interleukin 6)* | **F** | CTGCAAGAGACTTCCATCCAG |
| | **R** | AGTGGTATAGACAGGTCTGTTGG |
| *Lamp1 (Lysosomal-associated membrane protein 1)* | **F** | CAGCACTCTTTGAGGTGAAAAAC |
| | **R** | ACGATCTGAGAACCATTCGCA |
| *Mcoln1 (Mucolipin-1)* | **F** | TCATTGCACTCATCACCGGC |
| | **R** | CCAGATGTGGGGCTATCCTG |
| *Mcp1 (Monocyte Chemoattractant Protein 1)* | **F** | TTAAAAACCTGGATCGGAACCAA |
| | **R** | GCATTAGCTTCAGATTTACGGGT |
| *Mcp3(Monocyte Chemoattractant Protein 3)* | **F** | GCTGCTTTCAGCATCCAAGTG |
| | **R** | CCAGGGACACCGACTACTG |
| *P62 (SQSTM1/Sequestosome 1)* | **F** | GCTGCCCTATACCCACATCT |
| | **R** | GGCCTTCATCCGAGAAAC |
| *Pgc1b (Peroxisome proliferator-activated receptor gamma coactivator 1 beta)* | **F** | TCCTGTAAAAGCCCGGAGTAT |
| | **R** | GCTCTGGTAGGGGCAGTGA |
| *RagA (Ras-related GTP binding A)* | **F** | CTCGGGCGCTGTTTTCTGA |
| | **R** | CATGGCTGTATTGGGCATCAC |
| *RagB (Ras-related GTP binding B)* | **F** | CTAGCCAGCGTGACAACATCT |
| | **R** | GCCTCAAGGCATGACTGATAGT |
| *Tbp (TATA-box binding protein)* | **F** | ACGGACAACTGCGTTGATTTT |
| | **R** | ACTTAGCTGGGAAGCCCAAC |
| *Tgfb (Transforming growth factor beta)* | **F** | CCACCTGCAAGACCATCGAC |
| | **R** | CTGGCGAGCCTTAGTTTGGAC |
| *Tnfa (Tumor necrosis factor alpha)* | **F** | CAGGCGGTGCCTATGTCTC |
| | **R** | CGATCACCCCGAAGTTCAGTAG |
| *Uvrag (UV radiation resistance associated gene)* | **F** | GCAGACCACGAGACAGTTGA |
| | **R** | CATCGTGACGTTGCACACAG |

### Table S2. Primer sequences used for qPCR.

| Primary antibodies | | |
|---|---|---|
| **Targeted Protein** | **Dilution** | **Catalog #, Manufacturer** |
| 4E-BP1 | 1:1000 | 9644S, Cell Signaling |
| β-catenin | 1:1000 | 8480P, Cell Signaling |
| ATG7 | 1:1000 | 8558S, Cell Signaling |
| Akt (pan) | 1:2000 | 4691L, Cell Signaling |
| GSK-3β | 1:1000 | 9315S, Cell Signaling |
| LC3A/B | 1:1000 | 4108S, Cell Signaling |
| SQSTM1/p62 | 1:2000 | 5114S, Cell Signaling |
| p70 S6 Kinase | 1:1000 | 2708S, Cell Signaling |
| Phospho-4E-BP1 (T37/46) | 1:500 | 2855S, Cell Signaling |
| Phospho-Akt (S473) | 1:1000 | 4058L, Cell Signaling |
| Phospho-GSK-3β (S9) | 1:1000 | 9323S, Cell Signaling |
| Phospho-p70 S6 Kinase (T389) | 1:500 | 9234S, Cell Signaling |
| Phospho-TFEB (S211) | 1:500 | PA5-114662, Invitrogen |
| Rag A | 1:1000 | 4357S, Cell Signaling |
| Rag B | 1:500 | 8150S, Cell Signaling |
| TFEB | 1:1000 | A303-673A, Bethyl Laboratories |
| Vinculin | 1:2000 | sc-73614, Santa Cruz |
| **Secondary antibodies** | | |
| **Targeted host** | **Dilution** | **Catalog #, Manufacturer** |
| HRP-conjugated Goat anti-Rabbit IgG | 1:5000 | AS014, Abclonal |
| HRP-conjugated Goat anti-Mouse IgG | 1:10000 | 31430, ThermoFisher |

**Table S3. Primary and secondary antibodies.**
